# The Co-Evolution of Jawed Vertebrates and Interferon Regulatory Factor 5 Generates Unique Inflammation and Innate Immunity

**DOI:** 10.1101/2024.02.19.581058

**Authors:** Vanessa Hubing, Avery Marquis, Chanasei Ziemann, Hideaki Moriyama, Etsuko N. Moriyama, Luwen Zhang

## Abstract

The emergence of jaws in early vertebrates introduced a novel feeding apparatus and potent oral defenses but also increased the risk of physical injury and pathogen exposure. Innate immunity and inflammation constitute the body’s first line of defense against invading microbes and tissue damage, aiming to eliminate threats and restore internal homeostasis. Interferon regulatory factor 5 (IRF5) plays a critical role in orchestrating innate immunity and inflammation by regulating the transcription of genes that encode type I interferons (IFNs) and pro-inflammatory cytokines. Despite this, the evolution of IRF5 has remained poorly understood. We have identified the IRF5 and IRF6 genes in cartilaginous fish, including sharks. As cartilaginous fish represent one of the oldest surviving jawed vertebrate lineages, the presence of these genes suggests the genes have ancient origins potentially dating back hundreds of millions of years to early jawed vertebrates. Furthermore, our analysis shows that IRF5 has conserved nuclear export sequences and phosphorylation sites for activation throughout evolution from cartilaginous fish to humans, indicating these regulatory elements evolved early and have been maintained across jawed vertebrates. Additionally, the shift in subcellular localization of IRF5 from nucleus to cytosol, and of other interferon related IRFs, aligns with functional enhancements of IRFs in innate immunity and the emergence of IFNs across jawed vertebrates. This analysis implies that the evolution of jaws may have driven the emergence of new IRF members, the expansion of their functions, and the development of a unique inflammation and innate immune system.

## Introduction

The innate immune system and inflammation serve as the first line of defense against pathogens, injury, and stress. The innate immune system uses physical barriers, chemical mediators, and phagocytic cells to quickly recognize and eliminate invading microbes. Upon injury or infection, an inflammatory response clears pathogens and damaged cells while orchestrating repair processes like re-epithelialization and new tissue growth. Beyond basic defense, these evolutionarily ancient responses play crucial roles in tissue development, homeostasis, and regeneration. Overall, the nonspecific but highly integrated mechanisms of the innate immune and inflammation system are indispensable for appropriate development, physiological functioning, and disease prevention in complex multicellular organisms.

The evolution of jaws was a major innovation in early vertebrates, enabling a transition from filter feeding to active predation of larger, motile prey. This allowed access to new food resources but also posed new challenges. Powerful jaws risked physical injuries. Consuming larger prey exposed vertebrates to more ingested pathogens harbored in the animals they now eat. These selective pressures drove the evolution of quicker, more robust inflammatory and innate immune defenses to meet the challenges of a predatory lifestyle.

IFN regulatory factors (IRFs) are a small family of transcription factors with a variety of functions. IRF proteins share extensive similarity in the DNA-binding domain (DBD) located in the N-terminus, which is characterized by five well-conserved tryptophans (Ws in Supplemental Figure 1). The DBD region contains a helix-turn-helix structure and recognizes a DNA sequence known as IFN-stimulated response elements (ISRE) (Darnell et al., 1994). The C-terminal portion of IRFs contains the IRF-association domain (IAD), which is variable and defines their specific biological functions (Supplemental Figure 1). The IRF family has a variety of functions including, but not limited to, apoptosis, oncogenesis, host defense, and viral latency (Honda et al., 2006; Honda et al., 2005; Tamura et al., 2008; Zhang and Pagano, 2001, 2002).

IRF5 is a critical transcription factor that regulates inflammatory and innate immune responses. IRF5 is expressed in immune cells such as macrophages, dendritic cells, and lymphocytes. Upon activation by pattern recognition receptors, IRF5 moves from cytoplasm into the nucleus and induces the expression of pro-inflammatory cytokines as well as type I interferons (IFN) (Barnes et al., 2004; Takaoka et al., 2005). Through its control of inflammatory cytokines and type I IFNs, IRF5 exerts widespread effects on multiple arms of innate and adaptive immunity (Eames et al., 2016; Savitsky et al., 2010). Dysregulation of IRF5 is associated with uncontrolled inflammation in autoimmune diseases like systemic lupus erythematosus (Eames et al., 2016; Graham et al., 2007; Savitsky et al., 2010).

IRF6 and IRF5 belong to the same IRF subfamily (Angeletti et al., 2020; Ozato et al., 2007). Unlike IRF5, IRF6 plays a critical role in the formation of the jaw, palate, and skeletal system during embryonic development (Richardson et al., 2009). Mutations in IRF6 can cause cleft lip and palate, as well as deformities of the face and skull (Kondo et al., 2002).

Albeit exciting findings have been made since its discovery, molecular evolutionary studies of IRF5 and 6 have been limited (Angeletti et al., 2020; Du et al., 2018; Huang et al., 2010; Nehyba et al., 2009; Qi et al., 2018). Therefore, in this study, we examined IRF5 and related IRF protein sequences from a diverse range of animals. We have established that jawed fish (sharks) represent the earliest extant group possessing IRF5/6 during vertebrate evolution. In addition, we found that the newly expanded IRF members in jawed animals have relocated from the nucleus to the cytoplasm, apparently to more readily detect and transmit signals from outside the cell. Furthermore, given IRF6’s involvement in jaw and skeletal development, and that placoderms (420-360 million years ago) were the first jawed animals, our findings suggest the evolution of IRFs related to type I IFN systems may date back to placoderms.

## Results

### IRF5, 6, 4, and 8 first emerged in jawed vertebrates

Due to limited knowledge of the molecular evolution of IRF5, we conducted systematic searches for IRF5 and related IRF protein sequences, identifying their memberships based on similarities and phylogenetic locations (see Supplement Table S1). Our focus centered on IRF evolution during vertebrate development, and we gathered potential IRF sequences from lampreys (jawless fish) and cartilaginous fishes (jawed fish). Our phylogenetic analysis among IRF proteins clearly revealed the presence of IRF4/5/6/8 in various cartilaginous fishes (sharks; Chondrichthyes). Despite being annotated as "IRF5-like," "IRF6-like," "IRF8-like," or "IRF4-like" in public databases, the "-like" annotation denotes similarity and potential evolutionary relationships to known genes, with uncertainty about precise orthology or paralogy. Therefore, our phylogenetic analysis clarified these relationships (see Figure 1; Supplemental Table S1).

**Figure 1:**
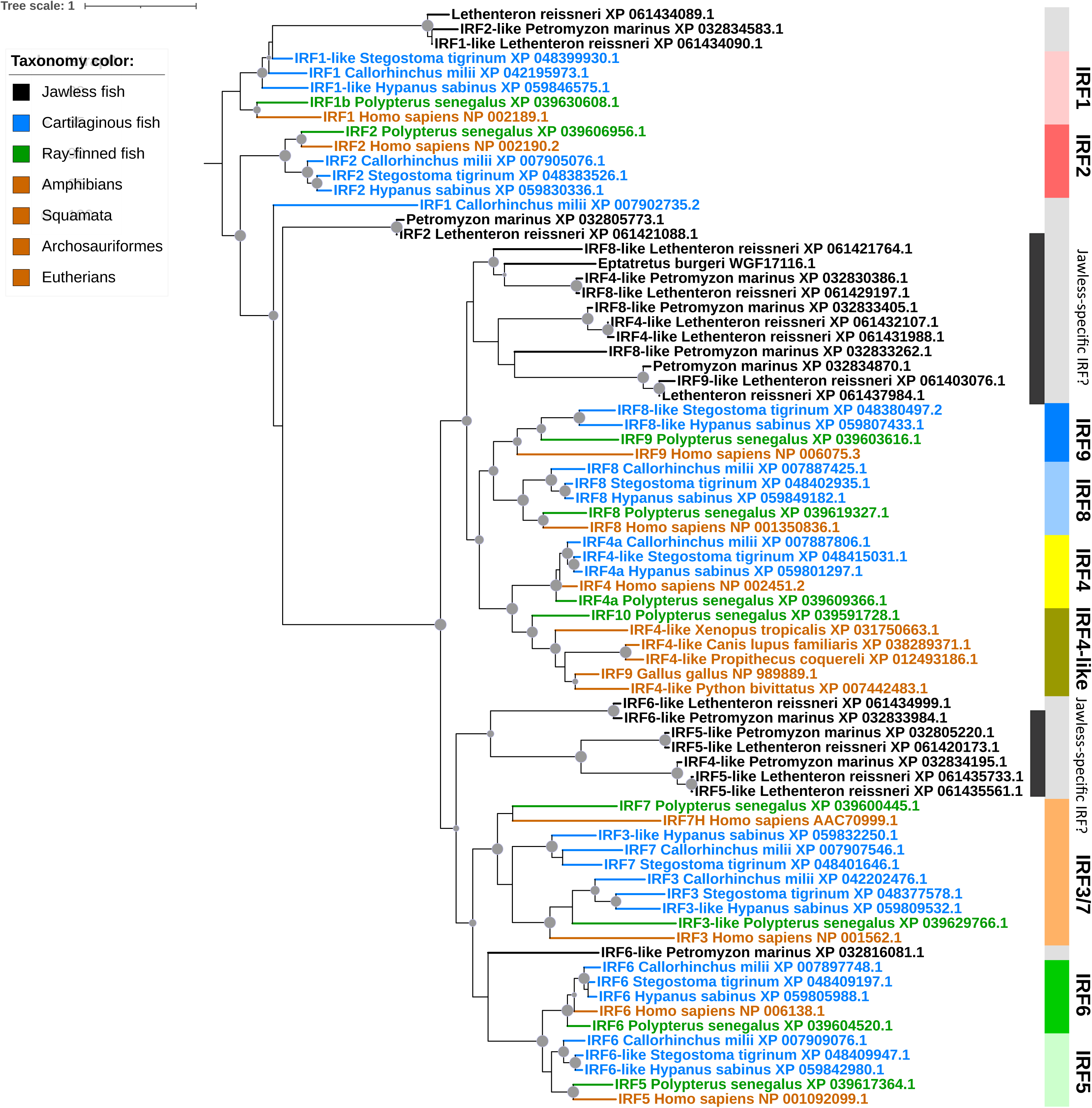
Phylogenetic relationships among IR5, IRF6, and IRF8 protein families. The maximum likelihood phylogeny was reconstructed using the IRF protein sequences from jawless and jawed animals from various species. The internal nodes supported by 80% or higher by both ultrabootstrap and SH-aLRT branch test are denoted by gray dots. The size of the dots is correlated with the ultrabootstrap values (80-100%). External branches and sequence names are colored based on the taxonomic groups as shown in the color legend. Supplementary Tables S1 for the protein sequences used.

Our data indicates that IRF5, 6, and surprisingly IRF4 and 8 first emerged in cartilaginous fishes, coinciding with the appearance of a primitive type I IFN system (Secombes and Zou, 2017). Considering various analyses, including ours and others (Angeletti et al., 2020; Drury et al., 2023; Huang et al., 2010; Kasamatsu et al., 2010; Nehyba et al., 2009), it appears that IRF3-9 have likely not been identified in Cephalochordata (lancelets), Urochordata (tunicates), Agnatha (lampreys), or other jawless vertebrates. In jawed vertebrates starting with cartilaginous fishes, IRF3-9 were present, leading to a significant expansion of the IRF family. Interestingly, jawless fish appear to possess distinct groups of IRFs (Figure 1, black bars on the right). Further research is necessary to classify and understand the functions of these unique IRFs.

### IRF5 has conserved nuclear export sequences and activation domains during evolution

The nuclear export sequences (NES) and activation domain (AD) for phosphorylation are two regions in IRF5 essential for its functions (Supplemental Figure 1). We compared the NES and AD sequence regions among many known IRF5 proteins and found that both sequences are conserved during evolution (Figure 2; Supplemental Table S2). In primitive sharks, IRF5 NESs are already established as functional components of the protein. As shown in Figure 2, the conserved NES sequences fit the consensus NES sequences defined in previous studies (la Cour et al., 2004; Xu et al., 2012). In addition, the AD sequences show similar evolutionary conservation. Furthermore, we analyzed putative structures of shark IRF5 and did a structural comparison between human and shark IRF5. As shown in Supplemental Figure 2, both NES and AD sequences are structurally similar and presumed to be involved in controlling the arrangement of molecules. Our data suggest that key functional domains governing IRF5 activity arose early in jawed vertebrates and have been maintained under purifying selection, highlighting their critical roles in regulating IRF5-mediated inflammation and antiviral immunity throughout vertebrate evolution.

**Figure 2.**
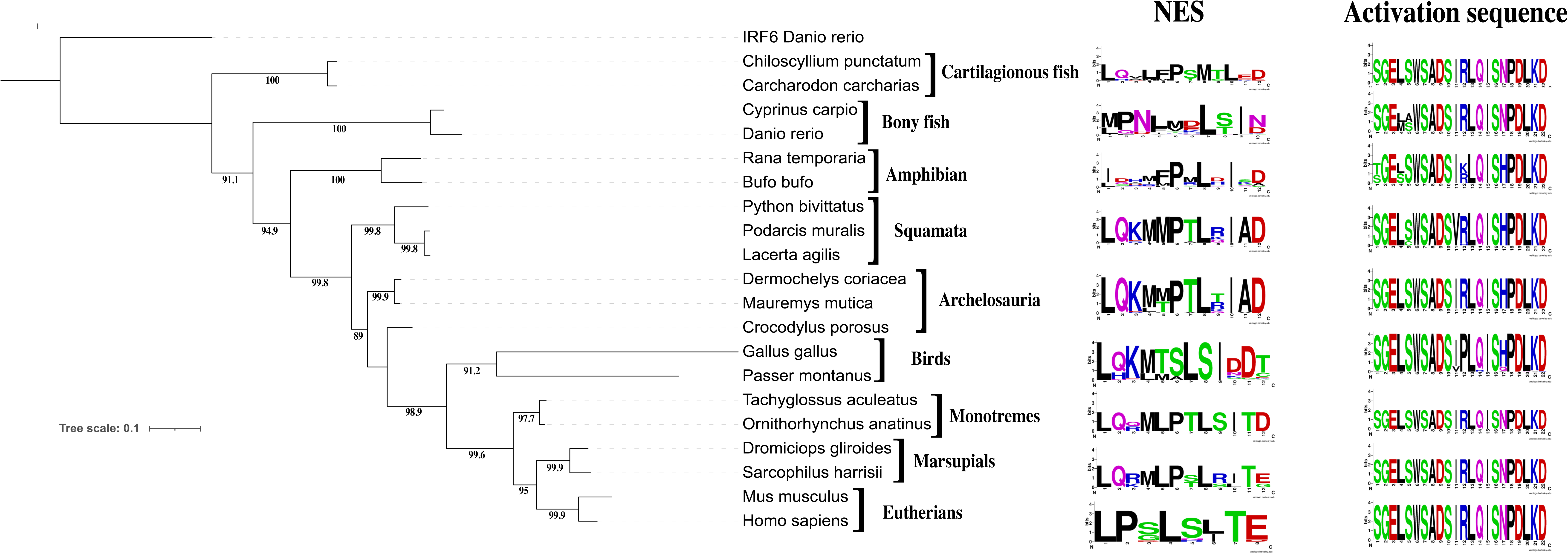
Conserved amino acid sequences found in the primitive IRF5 proteins in jawed animals. A. The maximum likelihood phylogeny was reconstructed using the IRF protein sequences from various species. The internal nodes supported by 80% or higher by both ultrabootstrap and SH-aLRT branch test are denoted by gray dots. B. Sequence logos are used to illustrate the amount of sequence conservation for each position. The overall height of the stack of letters indicates the sequence conservation at each position. The height of symbols within each stack indicates the relative frequency of each amino acid. The multiple sequence alignments were generated using protein sequences listed in Supplementary Table 2. The illustrated regions correspond to aa 150-161 (NES) and aa 447-468 (AD sequences) in the human IRF5 (NP_001092099.1).

### STAT2 first appeared in Jawed Animals

In IFN signaling, the signal transducer and activator of transcription-2 (STAT2) interacts with other proteins, STAT1 and IRF9, to form an IFN-stimulated gene factor 3 (ISGF3) complex that then translocates into the nucleus, where it binds specific DNA sequences and directs production of interferon-stimulated genes (ISGs) (Lau et al., 2000). IRF9 constitutively binds STAT2 in the cytoplasm without stimulation, and this binding is necessary for shuttling between the nucleus and cytoplasm (Banninger and Reich, 2004; Fink and Grandvaux, 2013; Rengachari et al., 2018). IRF9 lacks a NES but possesses a nuclear localization signal (NLS) within its DBD (Lau et al., 2000). Conversely, STAT2 has a functional NES but lacks an NLS. Therefore, without STAT2, IRF9 localizes to the nucleus (Lau et al., 2000; Paul et al., 2018).

To study the importance of IRF9 cytoplasm localization, we examined STAT2 evolution in jawless and jawed vertebrates. We gathered potential STAT sequences from lampreys (jawless fish) and cartilaginous fishes (jawed fish). Phylogenetic analysis revealed the presence of STAT2 in cartilaginous fishes (sharks; Chondrichthyes), while many “STAT-like” sequences in databases denote uncertainty about precise orthology. Our analysis clarified these evolutionary relationships (Figure 3; Supplemental Table S3). The presence of STAT2 in sharks suggests IRF9 was cytoplasmic early in IFN system evolution.

**Figure 3:**
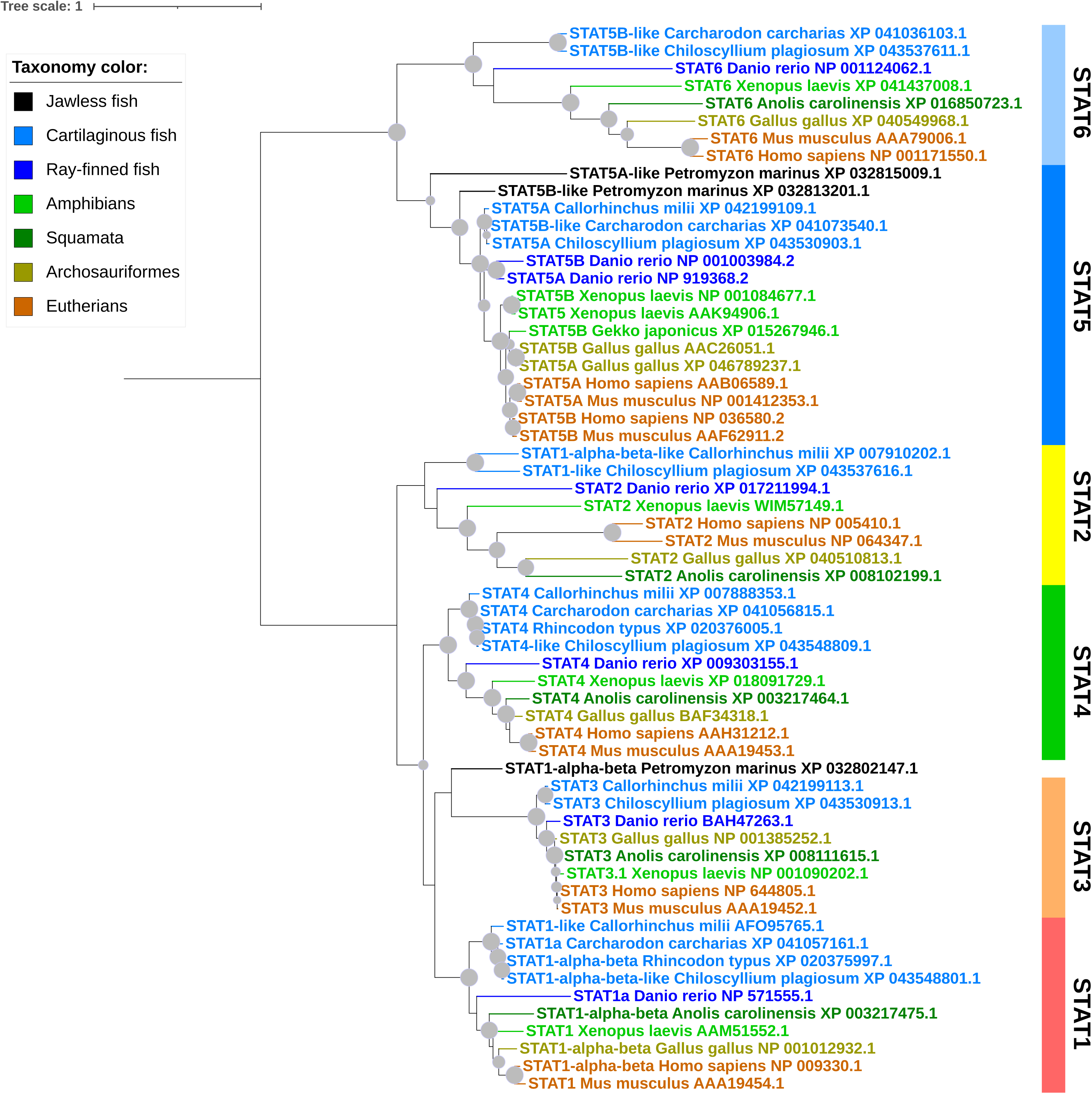
Phylogenetic relationships among STAT2 protein families. The maximum likelihood phylogeny was reconstructed using the STAT protein sequences from jawless and jawed animals from various species. The internal nodes supported by 80% or higher by both ultrabootstrap and SH-aLRT branch test are denoted by gray dots. The size of the dots is correlated with the ultrabootstrap values (80-100%). External branches and sequence names are colored based on the taxonomic groups as shown in the color legend. Supplementary Tables S3 for the protein sequences used.

### Jawed Animal-Specific IRFs Exhibit Nuclear-to-Cytoplasmic Relocation

Under basal conditions, IRF5 is predominantly localized in the cytoplasm, where it remains inactive. Upon activation by various stimuli, including microbial components and danger signals, IRF5 undergoes nuclear translocation, enabling its interaction with DNA and subsequent transcriptional regulation of target genes involved in immune responses. The cytoplasmic localization of IRF5 serves as a regulatory mechanism, preventing its constitutive activation and ensuring a controlled immune response. This dynamic subcellular localization of IRF5 is essential for its proper function in regulating innate immunity and inflammation. Similar mechanisms are also present for IRF3 and IRF7 (**Figure 4**)

**Figure 4:**
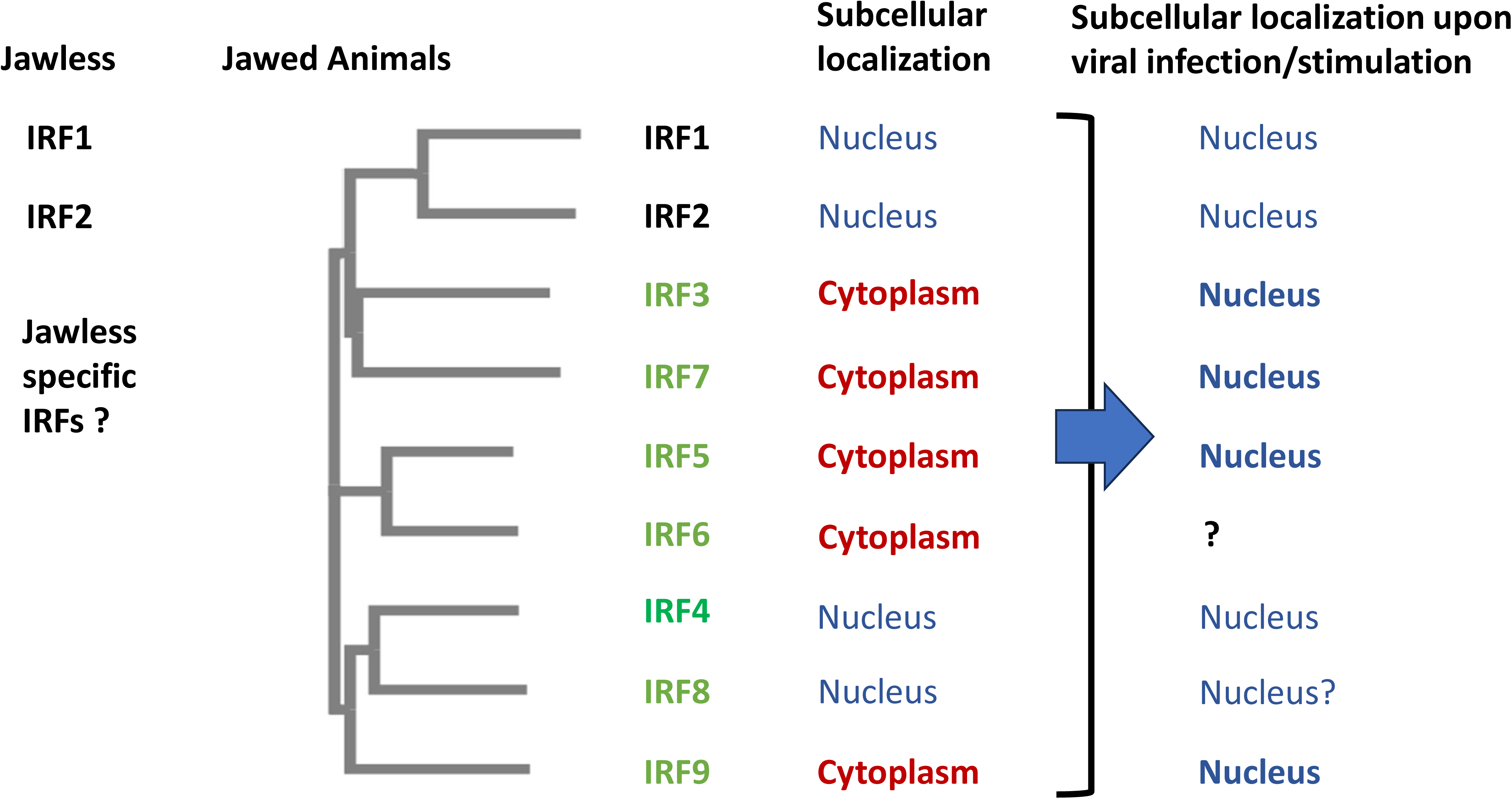
Comparisons between the IRF members in Jawless and Jawed Animals. **Left first column:** Jawless IRF members. **Left second column:** Phylogenetic tree of human IRFs. The newly expanded members are listed in green font. **Third column from the left:** Subcellular localization of those IRF members in the unstimulated state. **Last column (rightmost column):** Subcellular localization of those IRF members in the stimulated state, such as during viral infections.

It is known that IRF3/7 are cytoplasmic proteins that translocate into the nucleus upon viral infection (Glanz et al., 2021; Hiscott et al., 1999; Ning et al., 2011; Zhang and Pagano, 2001, 2002). Because STAT2 is necessary for cytoplasm retention of IRF9, the presence of STAT2 in sharks suggests IRF9 was cytoplasmic early in IFN system evolution (Figure 3).

IRF8 is the closest relative to IRF9 and IRF4 (Angeletti et al., 2020). Unlike the predominant cytoplasmic presence of IRF9, IRF8 is mainly nuclear (Laricchia-Robbio et al., 2005; Minderman et al., 2017; Schönheit et al., 2013). IRF8 apparently lacks physical interaction with STAT2, which may explain its nuclear localization. IRF1,2,4 are also well-known nuclear proteins (Schaper et al., 1998; Vora et al., 2021).

Therefore, with the emergence of jawed vertebrates, five newly expanded IRF members - IRF3/5/67/9 - are localized in the cytoplasm without stimulation, while IRF4/8 are nuclear. IRF3/5/7/9 are IFN pathway related IRFs. In summary, a nuclear-to-cytoplasmic shift accompanied the expansion and diversification of IRFs in jawed vertebrates (Figure 4).

### Birds have unique signature sequences for IRF5

While the overall inflammation and innate immunity systems are similar, birds and mammals differ significantly in the precise composition, diversity, and regulation of their innate immune and inflammatory mechanisms. Millions of years of evolution led to unique specializations in the bird lineage. Previously we compared the DNA-binding domain (DBD) regions among nine IRF families and identified YDG as a signature sequence for the IRF5/6 subfamily (Angeletti et al., 2020). By comparing the bird protein sequences within the IRF5/6 subfamily to those in mammals, the conserved signature motif for IRF5 (YDG) in birds is preferentially changed to other sequences, such as FDG and VDG (Figure 5; Supplemental Table S2).

**Figure 5:**
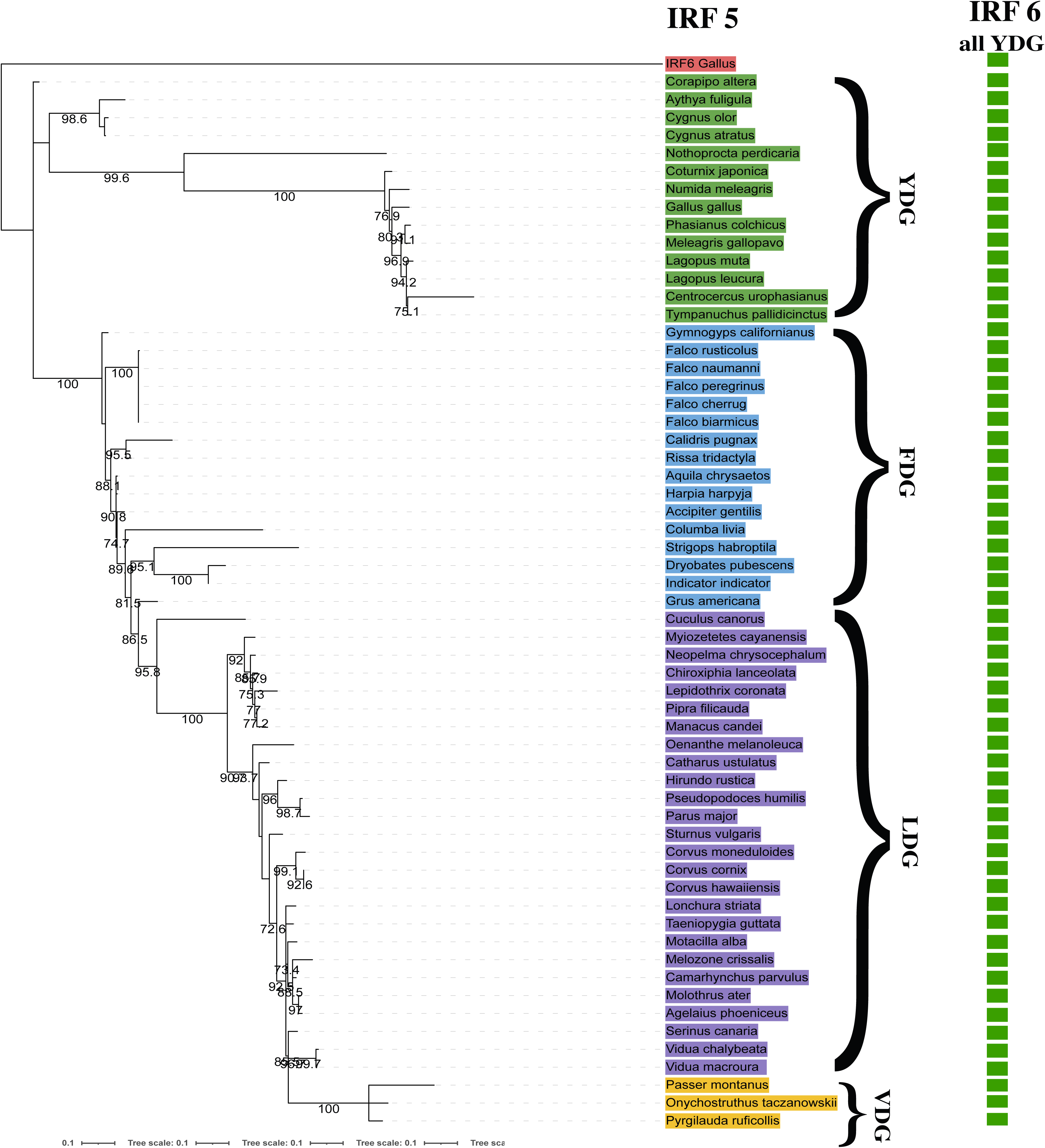
Distribution of 5 YDG motifs in birds. The maximum-likelihood phylogeny of IRF5 proteins is shown with the bootstrap supporting values (%) at the nodes where the supporting values are equal to or higher than 70. IRF6 sequences are used as the outgroups. Colors represent changes. See Supplementary Table S2 for the protein sequences used.

Clearly the conserved motif YDG is favored through vertebrate evolution. Since the DBD region of IRF proteins is generally more conserved, it suggests the important functional role of the YDG motif in IRF5/6 proteins in mammalian lineages, where it must have been under strong evolutionary constraints. Why does the bird lineage change the conserved YDG motif to other FDG or VDG sequences? This substitution in birds suggests relaxation of functional constraints on the IRF5 DBD region compared to mammals, allowing greater sequence flexibility. Further studies are needed to determine if the variant bird motifs lead to differences in IRF5 DNA-binding specificity or activity. The IRF5 DBD changes may represent an avian-specific adaptation impacting transcriptional networks controlling inflammation and antiviral immunity.

## Discussion

### IRFs are Evolving from Nuclear Transcription Factors to Cytoplasmic Signaling Molecules

Sharks, while not the earliest jawed vertebrates in the fossil record, have survived to the present day as the most ancient, jawed vertebrate group. They provide a living glimpse into the distant past of some of the earliest jawed fish. The primitive IFN system is likely to have developed in sharks (Secombes and Zou, 2017). By examining the evolution of IRFs in jawless and cartilaginous fish, the evolution of innate immunity has emerged.

Firstly, the IRF family has expanded its members. IRF3, 5, 7, and 9 first appeared during jaw evolution. All four are associated with IFN signaling: IRF3, 5, and 7 are involved in IFN induction, and IRF9 is a major factor in IFN signaling. Secondly, the IRF family has expanded its subcellular localization, from predominantly nuclear to cytoplasmic (Figure 4). Therefore, a major revolutionary jump in innate immunity was established during jaw development.

The evolution of IRFs to move from the nucleus to the cytoplasm has conferred several significant biological advantages in terms of evolutionary adaptation. Firstly, a cytoplasmic location allows for more rapid sensing and response to pathogens or cell damage, bypassing the need for signals to reach the nucleus. Cytoplasmic IRFs may directly interact with cytoplasmic pathogen components and danger signals, enabling autonomous activation of immune defenses. Secondly, cytoplasmic localization facilitates new interactions, modifications, and partnerships, potentially conferring broader innate sensing abilities or expanding inflammatory signaling cascades. This is particularly relevant for IRF9, which requires STAT2 to remain in the cytoplasm (Banninger and Reich, 2004; Fink and Grandvaux, 2013; Rengachari et al., 2018). Finally, by shuttling between the nucleus and cytoplasm, IRFs have expanded their functions from nuclear transcriptional factors to sensing and signal transduction molecules. IRF extrusion from the nucleus has likely conferred advantages in speed, magnitude, detection, regulation, interactions, or specialization, ultimately enhancing inflammation and innate immune responsiveness.

### The Evolution of the Vertebrate Jaw May Have Driven Revolutionary Expansions in IRF Membership and Function

The apparent distinct classes of IRF transcription factors in jawless fish versus jawed vertebrates provide molecular evidence that each lineage possesses unique immune systems and inflammatory responses. Jawless fish have a limited IRF family. In contrast, the IRF family expanded significantly in jawed vertebrates to include lineage-specific members like IRF3, IRF5, IRF6, and IRF8 (Figure 1). These additional IRFs are integral to specialized immune processes in jawed animals like IFN production. The apparent differences in IRF composition between jawless and jawed vertebrates strongly implicate major evolutionary divergences in antiviral defenses, adaptive immunity, and inflammation control.

The immune system and inflammation are intricate biological processes. Initially, these responses are triggered and regulated by external factors such as pathogens, toxins, and tissue injuries. In this sense, they operate reactively, responding to external influences rather than acting independently and proactively. Therefore, the association between newly expanded IRFs and jaw development strongly suggests that jaw formation functioned as a driving force in the development of the unique and robust type I IFN and inflammation system.

However, it’s worth noting that both the immune and inflammation systems possess memory, which enables them to continuously monitor for threats in a proactive manner after their initial encounters. In addition, newly evolved IRF6, although predominantly located in the cytoplasm, no evidence so far has linked the factor to immunity and inflammation. In humans, mutations in IRF6 cause two related orofacial clefting disorders - Van der Woude (VWS) and popliteal ptyergium syndromes (PPS) (Kondo et al., 2002; Zucchero et al., 2004). Mice deficient for Irf6 have abnormal jaw, limb and craniofacial development (Ingraham et al., 2006). Those data suggest that IRF6 itself may be involved in jaw development from an evolutionary perspective. Interestingly, placoderms, an extinct class of armored fish, are often considered the first jawed vertebrates (Brazeau and Friedman, 2015; Brazeau et al., 2023). They lived from 420 to 360 million years ago and were the dominant vertebrate predators of their time. Although placoderms’ jaws were quite different from modern jaws, their primary functions were very similar: both are used for feeding and fighting. It is tempting to speculate that IRF6 may have been involved in jaw formation in placoderms and furthermore, IRF5, its close relative, might have mediated inflammation and innate immunity. Primitive IFN and inflammation systems may have originated around 420 million years ago, with the appearance of placoderms.

## Materials and Methods

### Searching of IRF and STAT proteins

The protein sequences of the human IRF protein sequences were used as the queries (see Supplementary Tables S1, S2, and S3 for accession numbers) to perform protein similarity searches using BLASTP against the non-redundant protein database at NCBI with the default options. Sequences were collected from mammals, reptiles, birds, amphibians, and fishes. When more than one isoforms were available from the same species, one isoform that was most similar to those from other species was selected. Protein sequences that were partial and too short were excluded. In total, all sequences collected in this study are listed in Supplementary Tables S1, S2, and S3.

### Phylogenetic analysis of IRF and STAT proteins

Starting with the IRF protein sequences obtained from the BLASTP searches, alignment and phylogenetic analysis were iteratively performed. After identical sequences were removed and grouping of IRF protein families was established, the alignment was performed again using each IRF family. The final alignment including all IRF sequences was performed using each of the IRF-family specific alignment as the profile and using the “merge” alignment of MAFFT with the E-INS-i iterative refinement method. The visualization of the phylogenies was performed using Interactive tree of Life (ITOL) website (Letunic and Bork, 2021). For STAT2 and other sequences, the same procedures were used.

### Sequence logos

A sequence logo was generated from the protein sequence alignment using WebLogo v3 (https://weblogo.threeplusone.com/create.cgi). For simplicity, the composition adjustment was suppressed. The amino acid “chemistry” color scheme was chosen.

### Structural Comparison Studies

The five-given shark-derived sequences and human-derived IRF5 (XP_047276292.1) were compared by BLAST, and the one with the highest total score (XP_051889933.1) was used. A three-dimensional model was predicted for these sequences on ColabFold server v1.5.5 (AlphaFold2 using MMseqs2). The predicted model’s DNA binding domain and core domain matched the experimentally solved structures, 7o56 and 3dsh, respectively. To obtain the same arrangement in human-derived IRF5 and shark-derived sequences, shark-derived sequences were remodeled in User-template mode of SWISS-model.

## Acknowledgments

This study was supported by the Collaboration Initiative Grant from the University of Nebraska (LZ, EM, and HM). VH, AM, and CZ were partially supported by the UCARE program at UNL. We would like to thank Ryan Herman for his initial efforts in collecting the data.

---

**Supplemental Figure 1:**
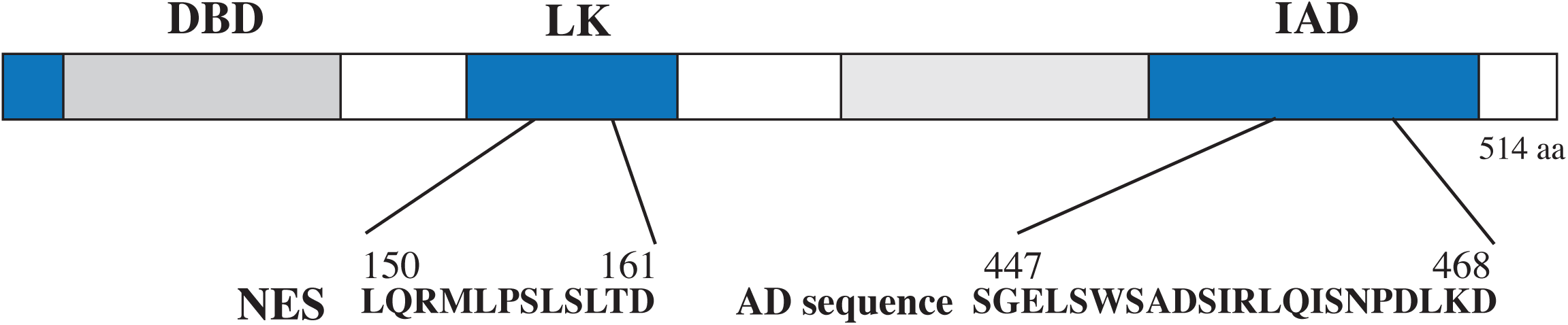
Domain structures of IRF5 protein. The DNA-binding domain (DBD), the IRF-association domain (IAD), the linker region (LK), activation domain (AD), five well conserved tryptophans (W) in DBD, and nuclear localization signal (NLS) and nuclear export signal (NES) are as shown. The illustrated regions correspond to aa 150-161 (NES) and aa 447-468 (AD sequences) in the human IRF5 (NP_001092099.1).

**Supplemental Figure S2:**
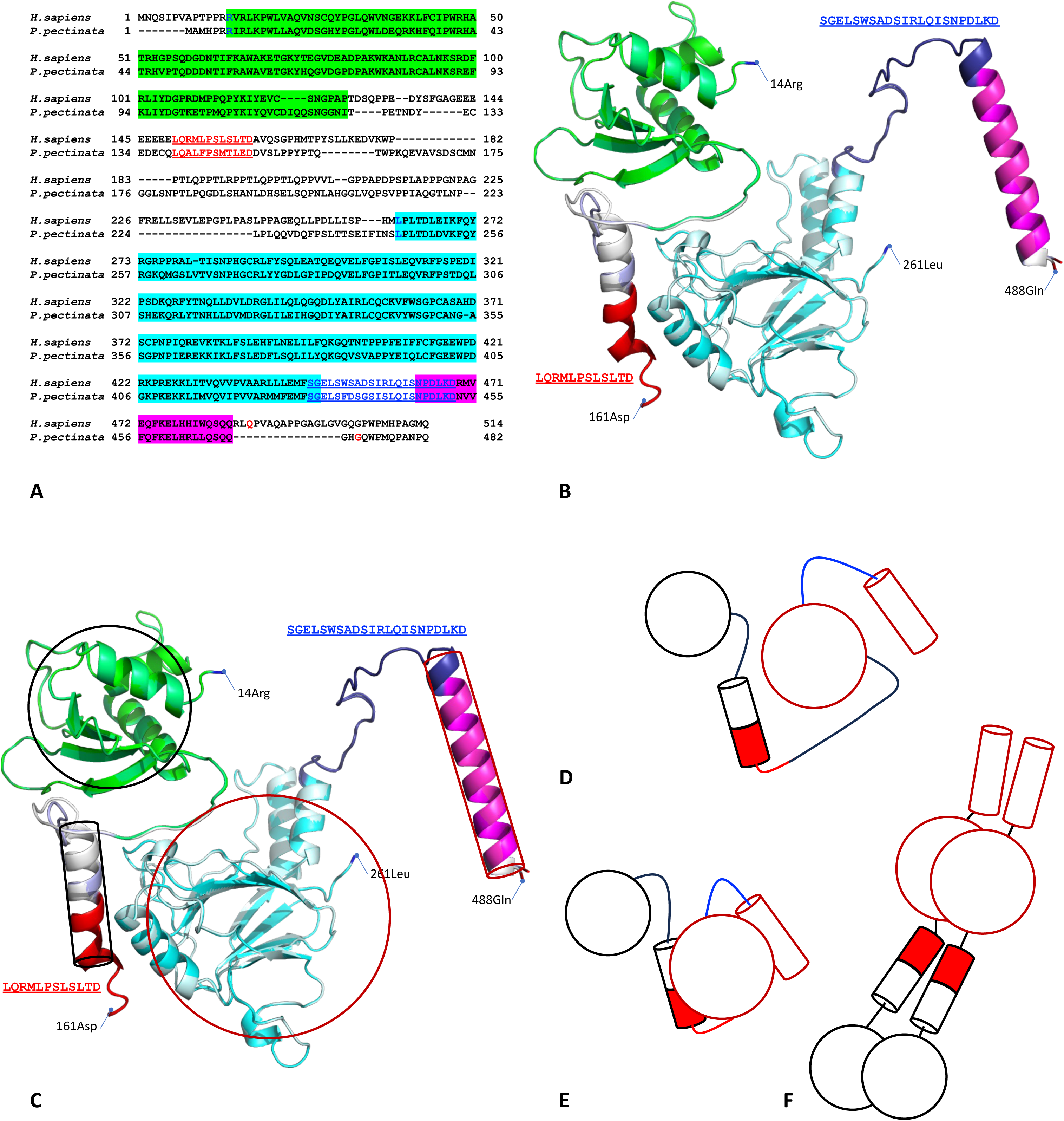
**A.** Human-derived (Homo sapiens) and shark-derived (Pristis pectinata) IRF5 sequences are used here. Highlighted sequences are structural domains, including the DNA-binding domain (green), core domain (cyan), and core domain outer helix (Helix 5 designated by core domain)(Chen et al., 2008). Sequences of interest are shown in red (LQRMLPSLSLTD in human) and blue (SGELSWSADSIRLQISNPDLKD in human). **B.** Superimposed structural model of IRF5 from human (light gray) and shark (light blue). Structural domains and color coding correspond to the sequences in Panel A. Amino terminus domains 14Arg–161Asp in human and carboxyl terminus domains 261Leu– 488Gln are shown. Internal loop connection amino carboxyl terminal are absent from the figure. **C.** Formation of Schematic Models DNA binding and core domains are shown as circles. Helices are shown as cylinders. **D.** A schematic model for IRF5 as modeled by AlphaFold2. **E.** Potential structure in the contracted autoinhibition mode. **F.** Potential structure in the DNA-binding mode in dimer format.

**Supplemental Table S1**: Sequences used to generate Figure 1.

**Supplemental Table S2**: Sequences used in this study to generate Figure 2.

**Supplemental Table S3**: Sequences used to generate Figure 3.

**Table S1.**
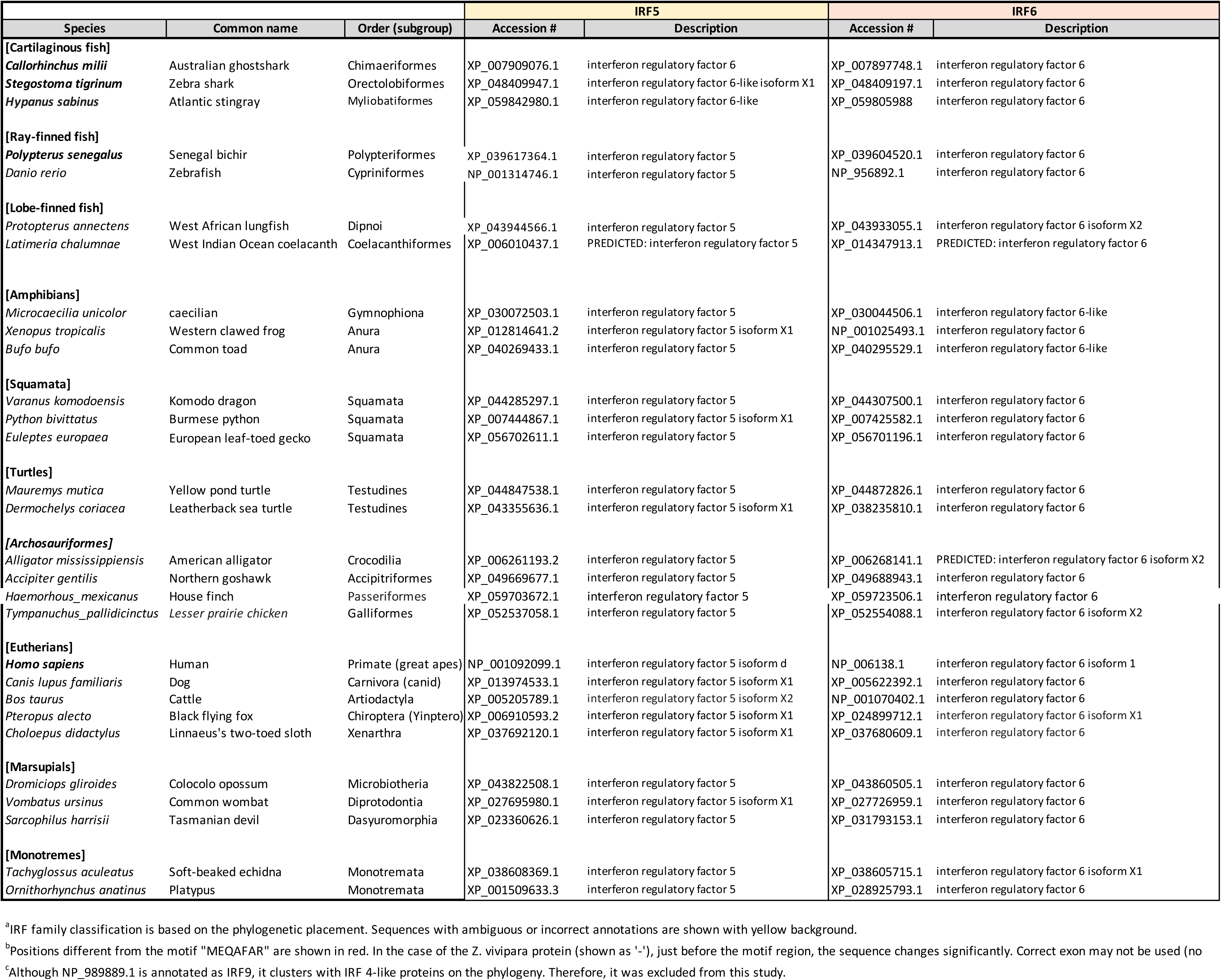
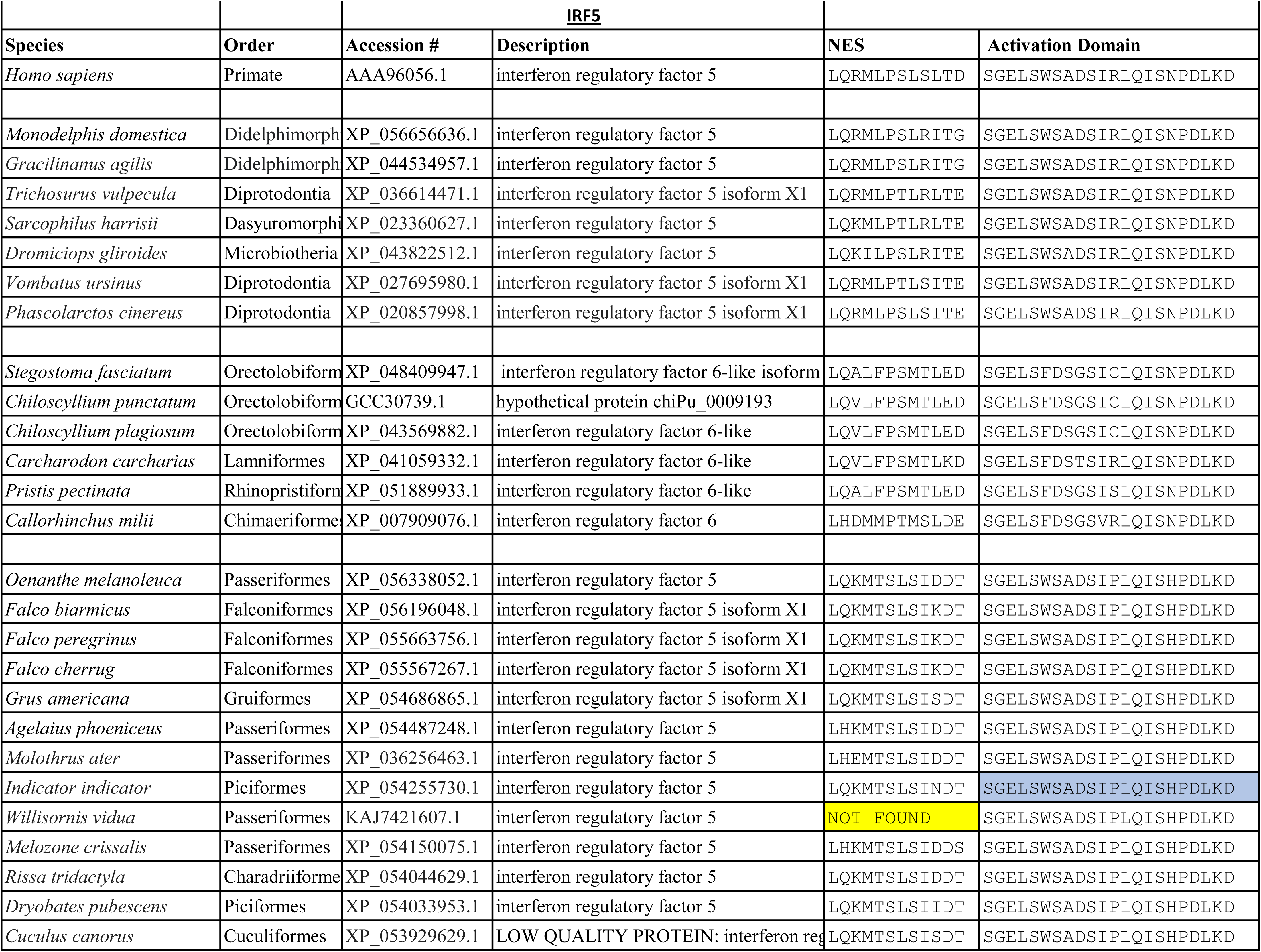
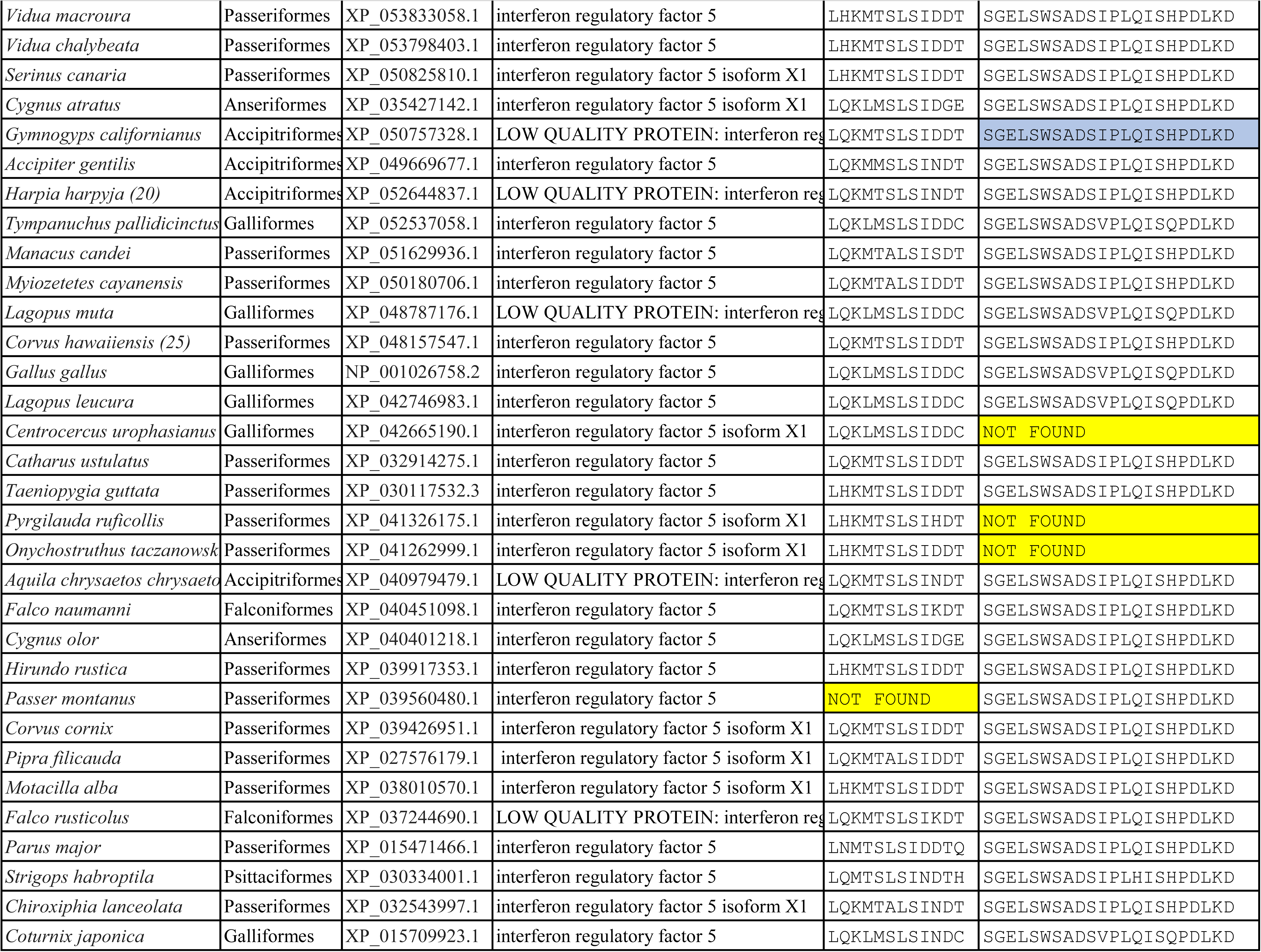
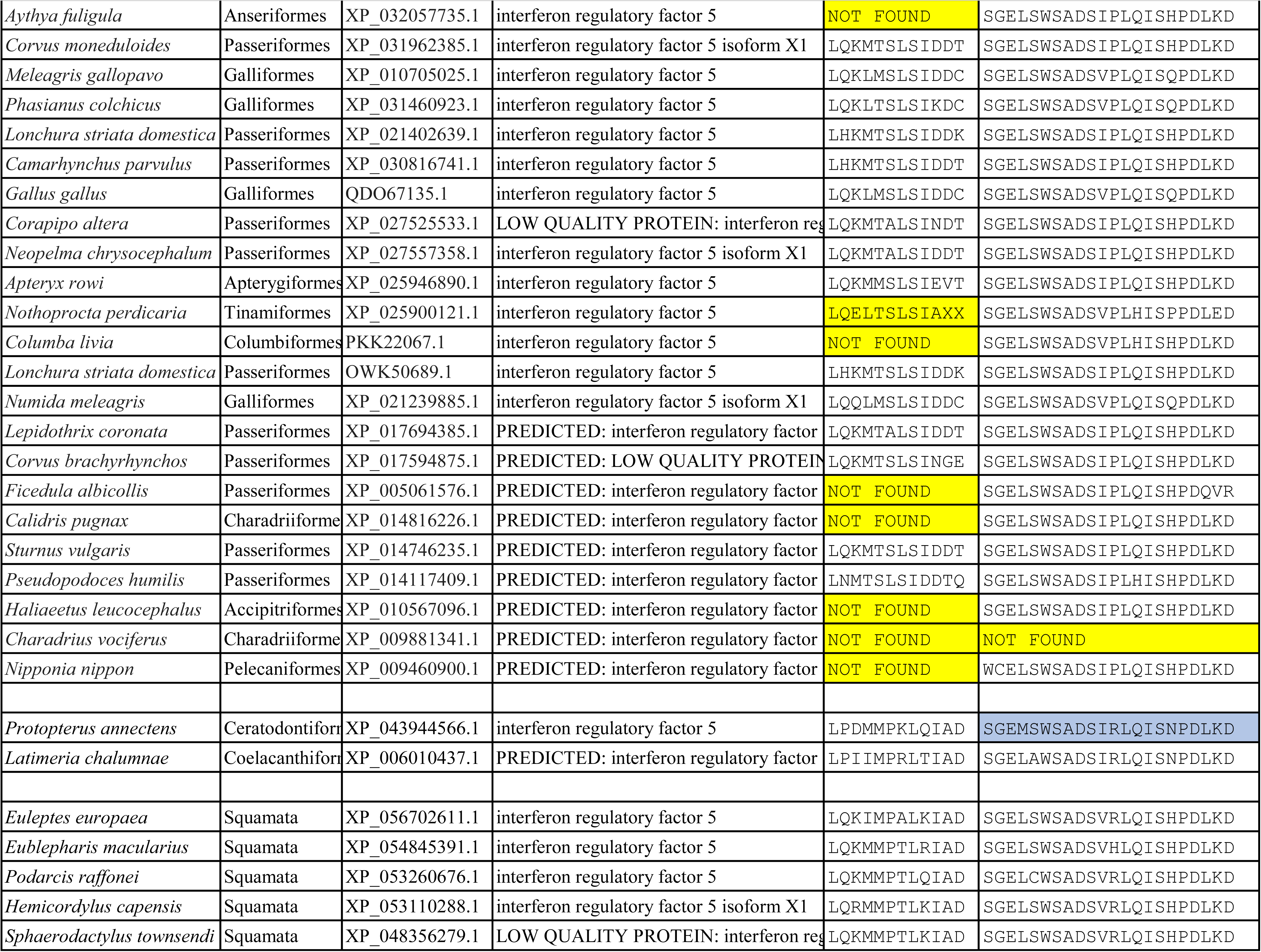
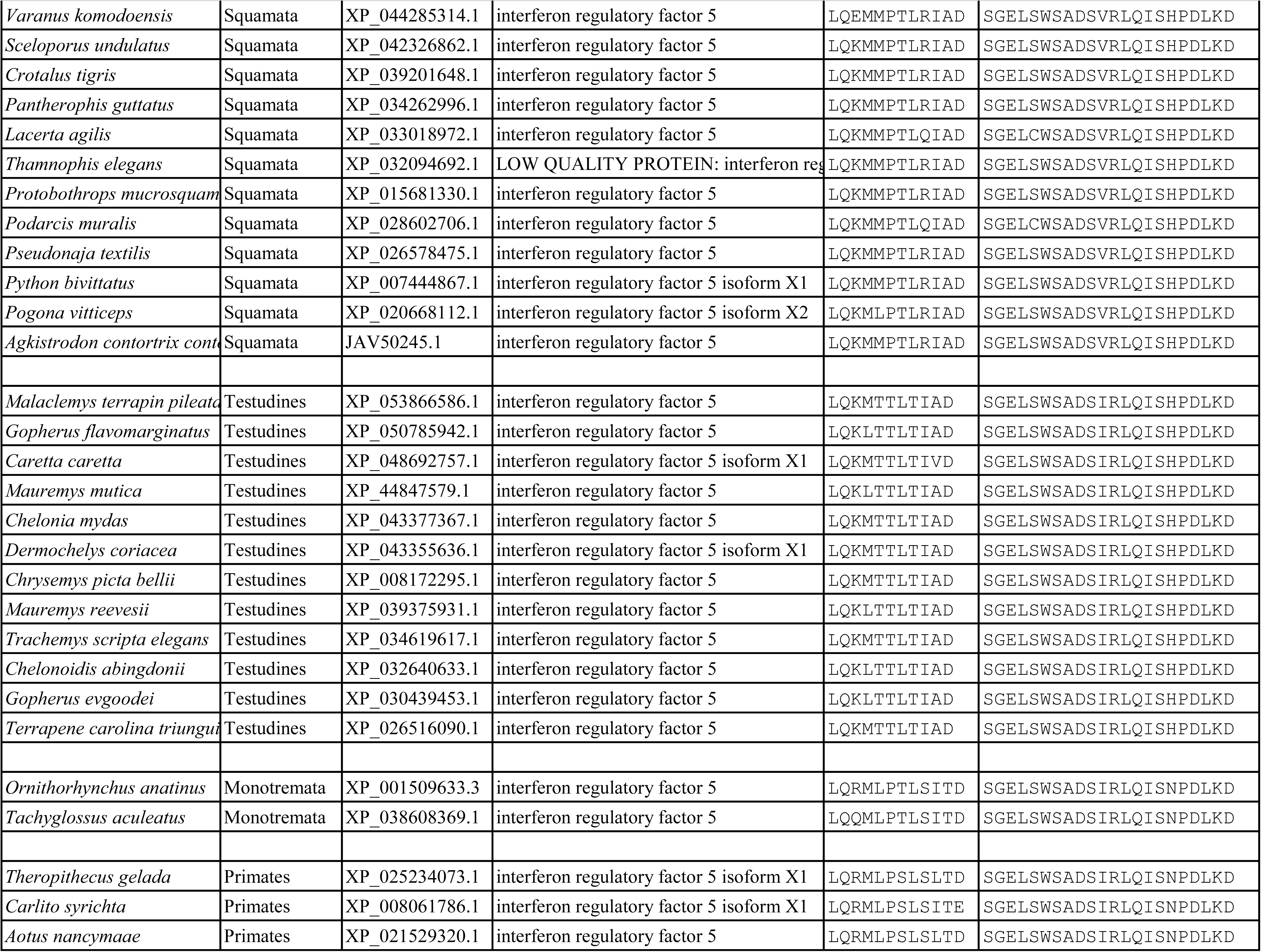
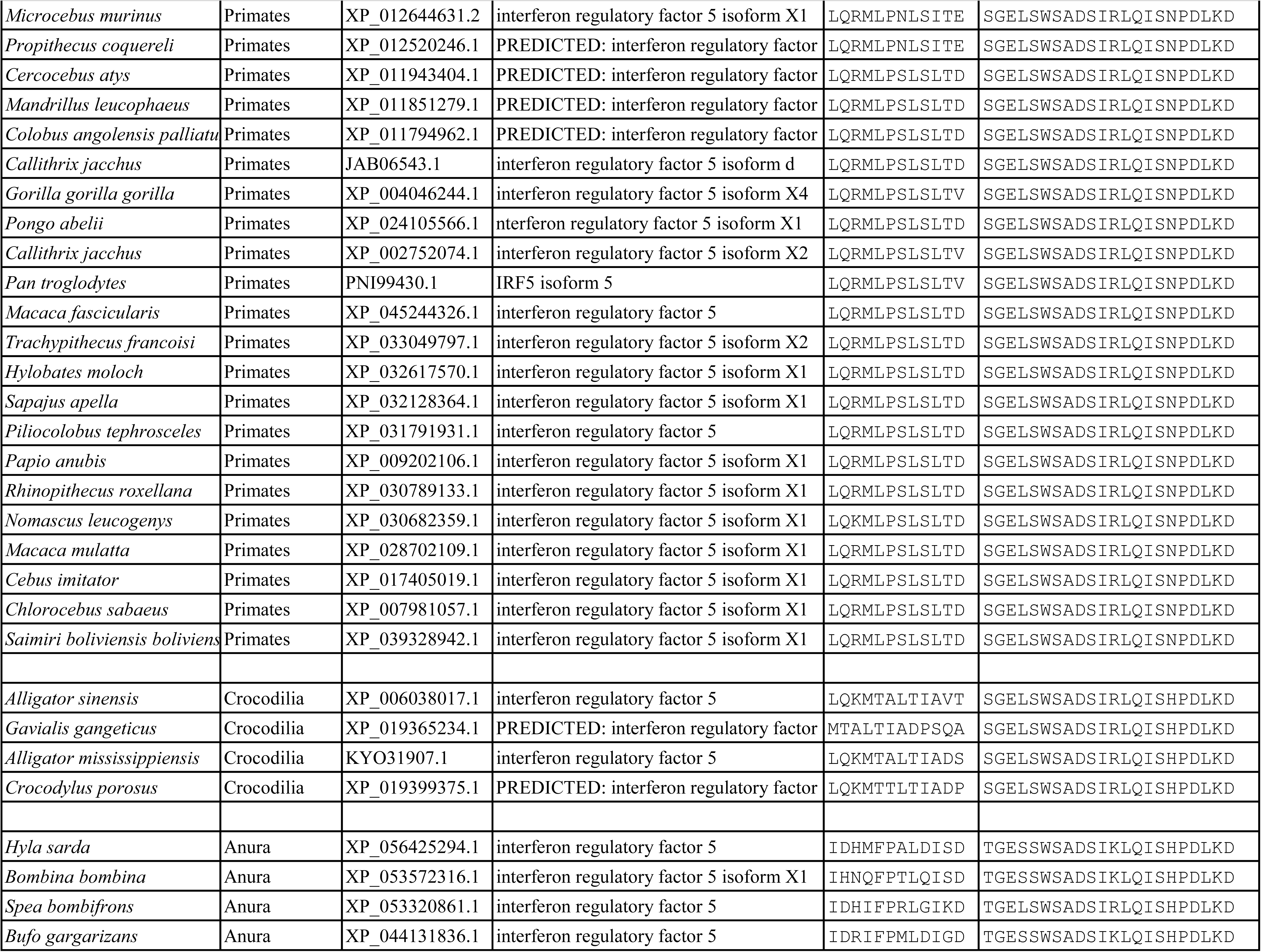
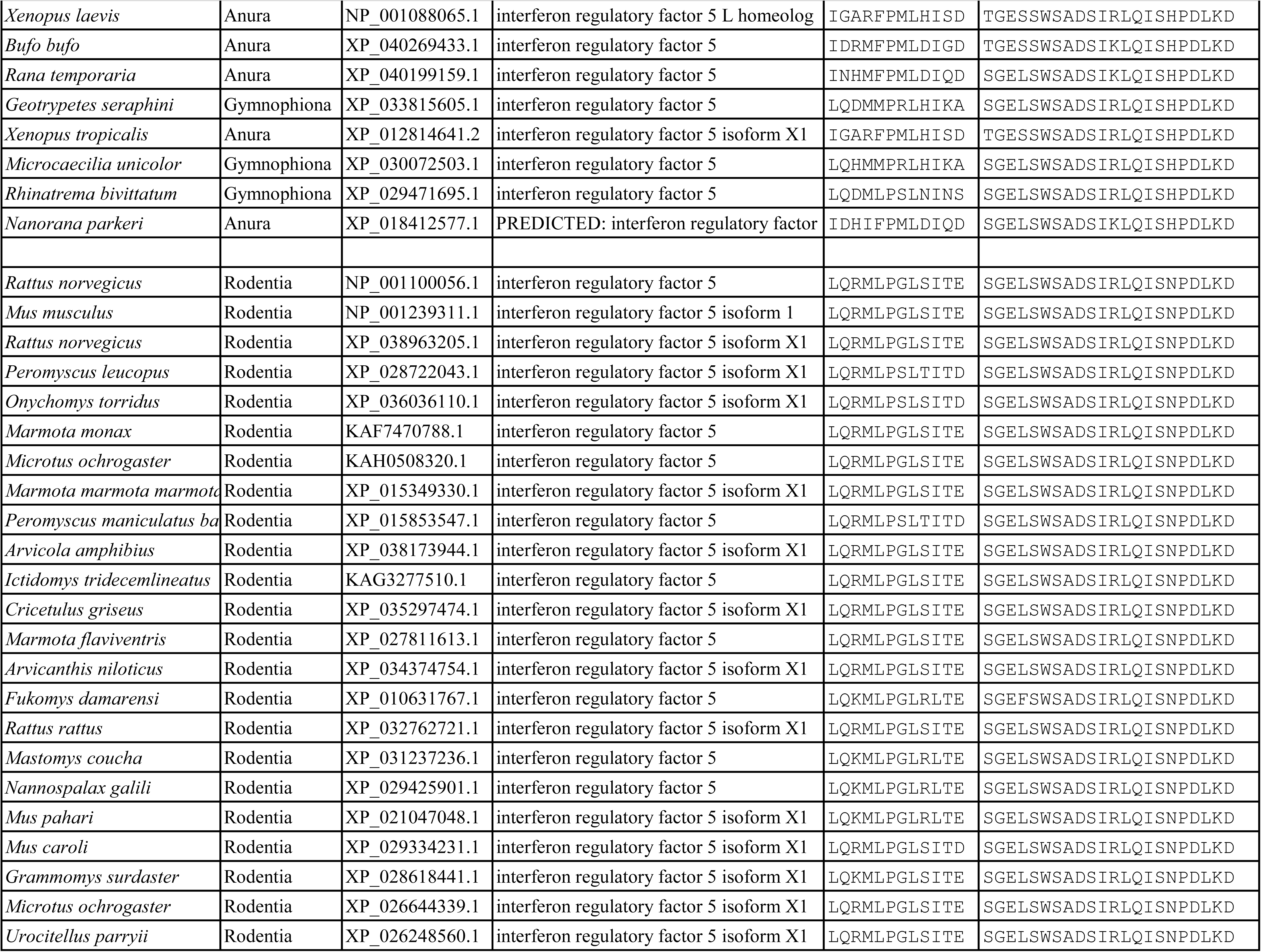
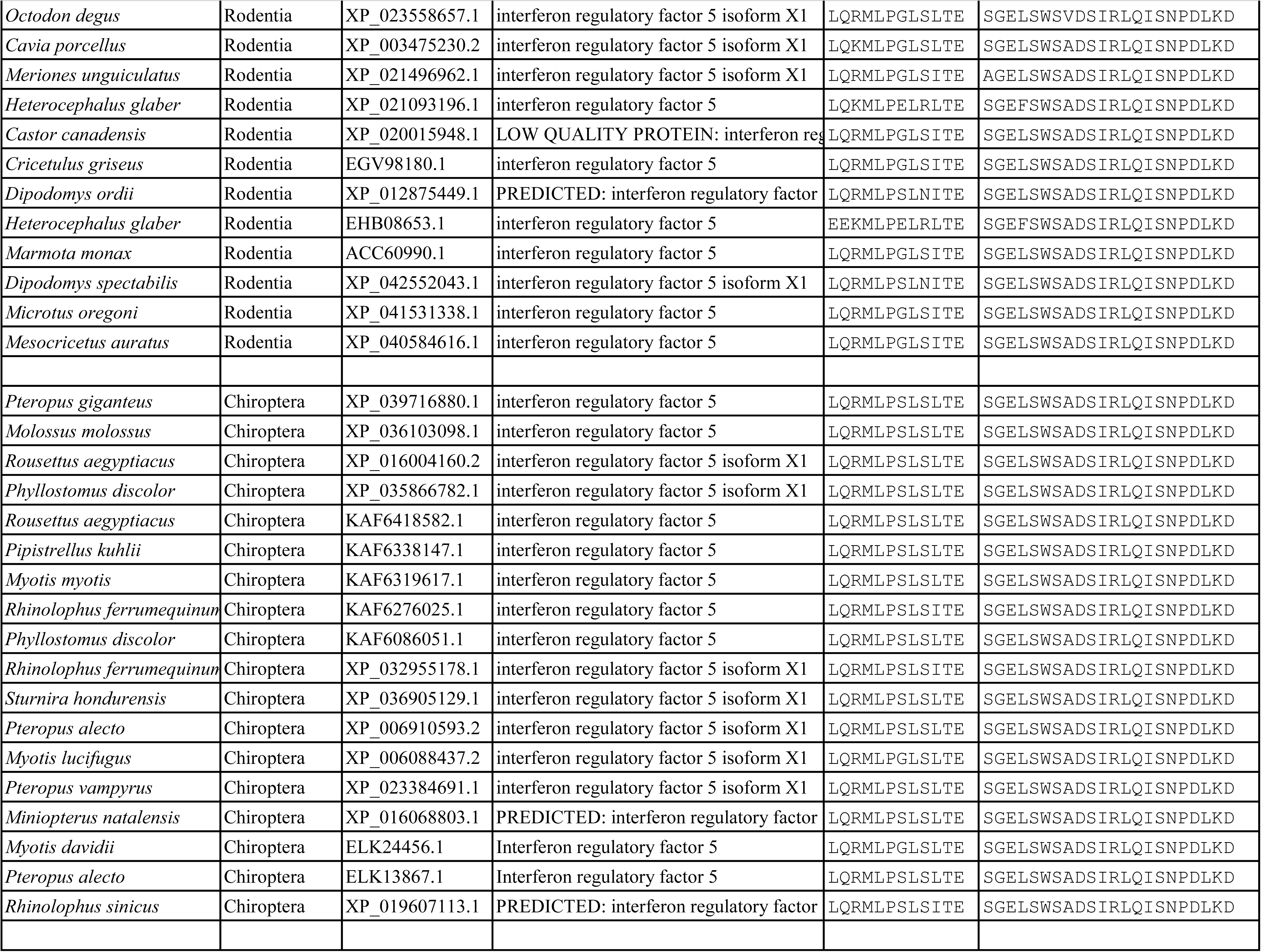
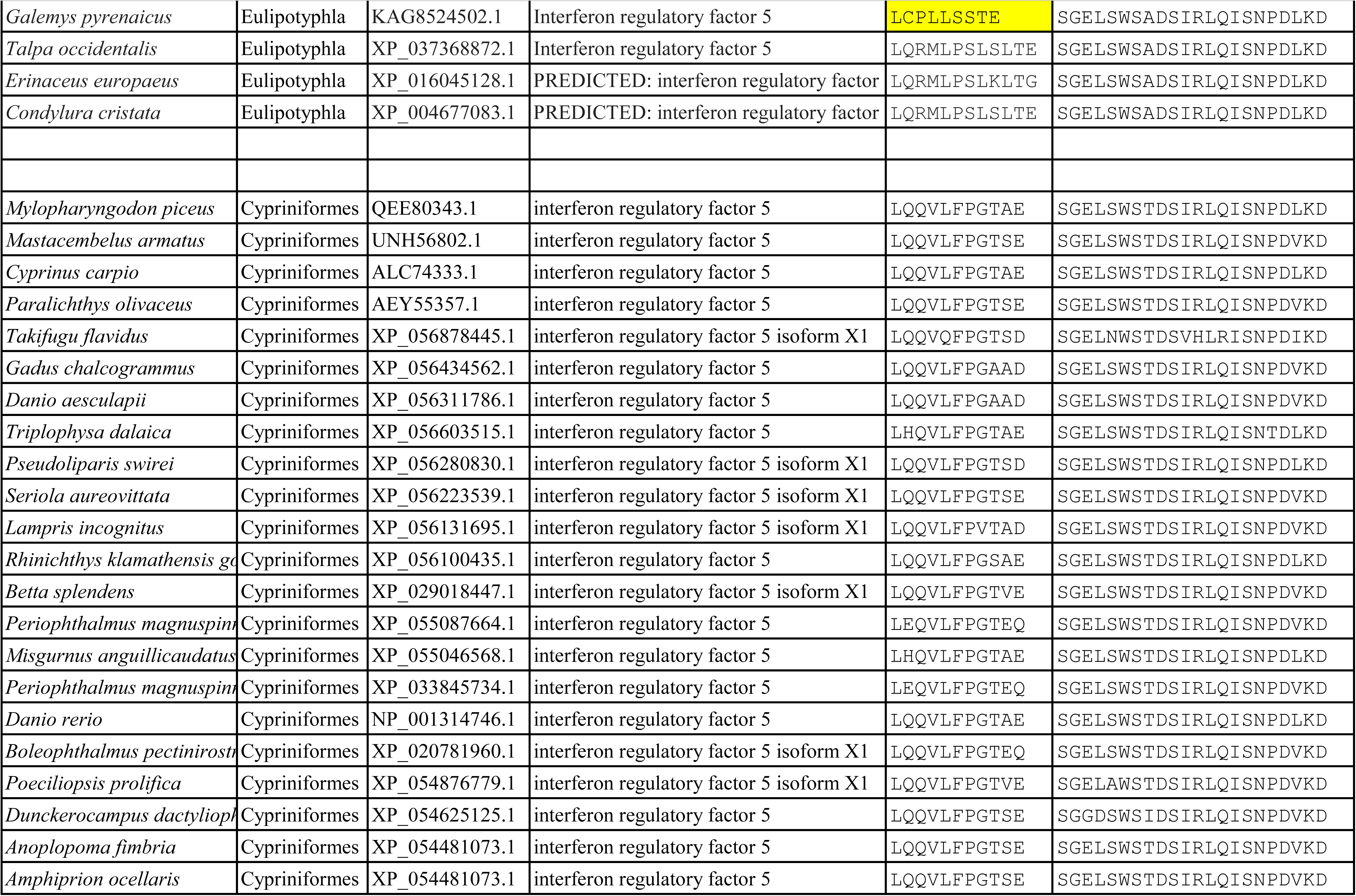
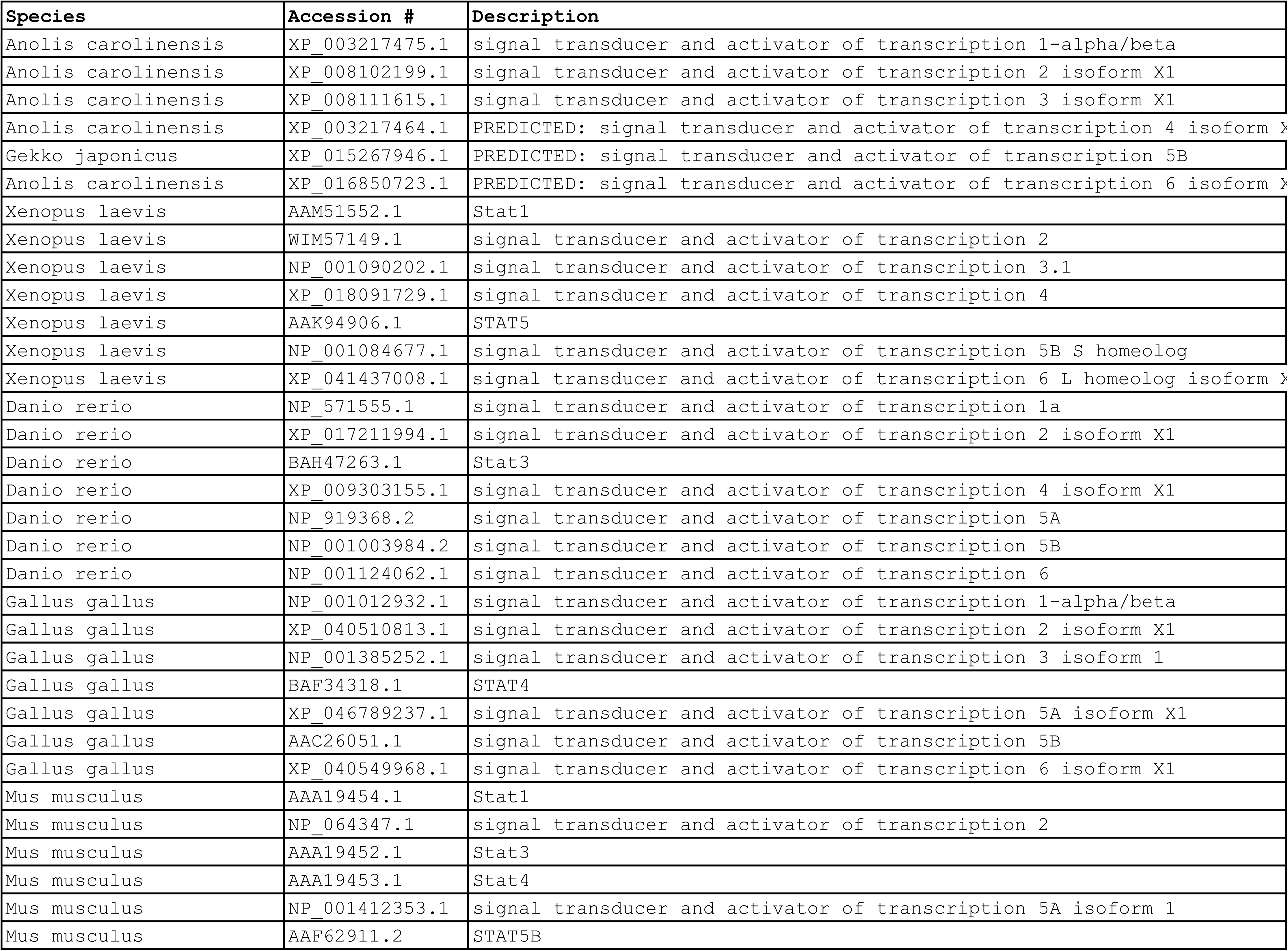
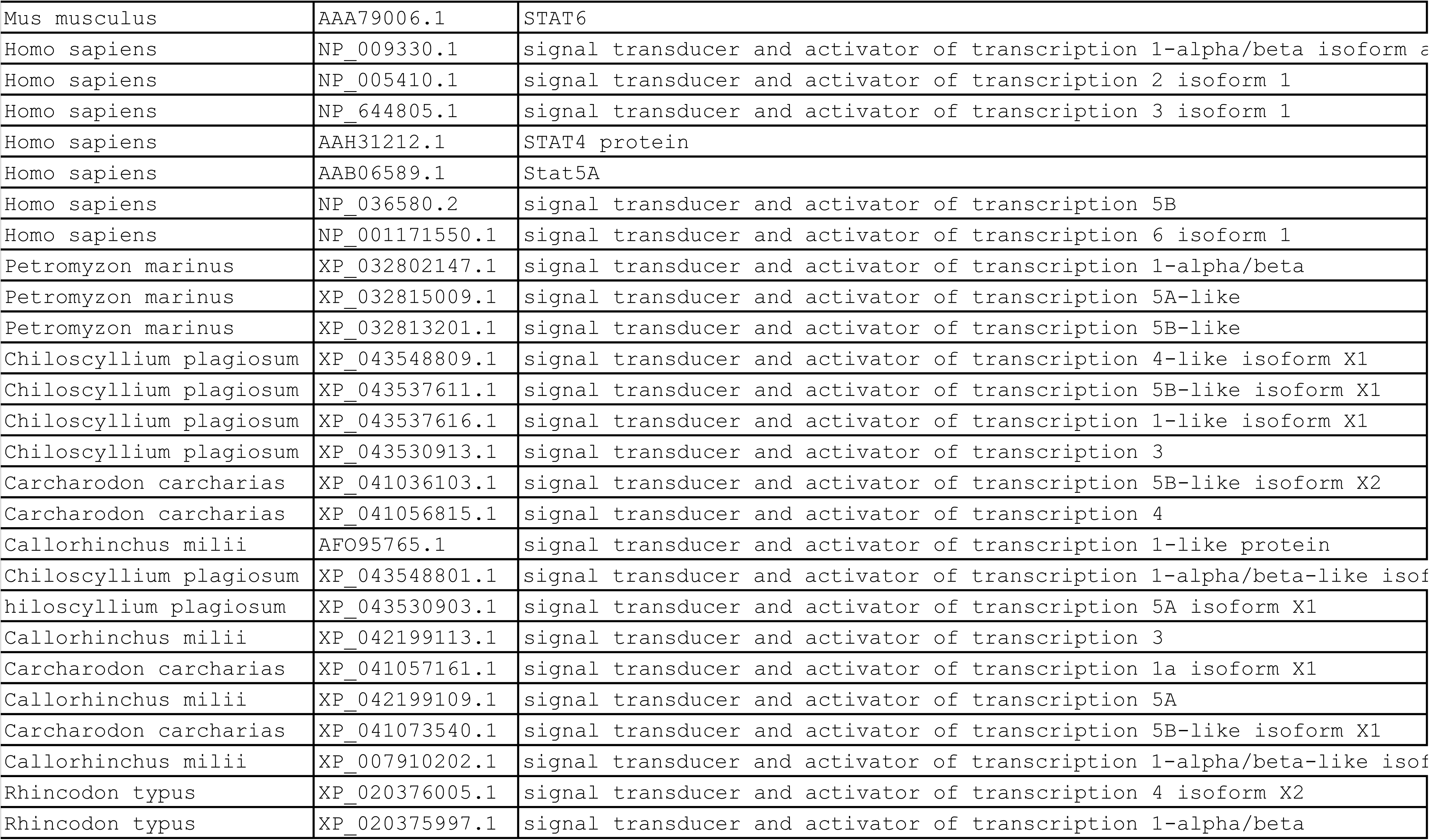
IRF5/6/4/8/9 potein sequences used in this study.^a^.

